# Recurrent gene amplification on Drosophila Y chromosomes suggests cryptic sex chromosome drive is common on young sex chromosomes

**DOI:** 10.1101/324368

**Authors:** Chris Ellison, Doris Bachtrog

## Abstract

Theory predicts that selfish genetic elements that increase their transmission are prone to originate on sex chromosomes but create strong selective pressure to evolve suppressors due to reduced fertility and distorted population sex ratios. Here we show that recurrent genetic conflict over sex chromosome transmission appears to be an important evolutionary force that has shaped gene content evolution of sex chromosomes in Drosophila. We demonstrate that convergent acquisition and amplification of spermatid expressed gene families are common on Drosophila sex chromosomes, and especially on recently formed ones, and harbor characteristics typical of meiotic drivers. We carefully characterize one putative novel cryptic sex chromosome distortion system that arose independently several times in members of the *Drosophila obscura* group. Co-amplification of the *S-Lap1*/*GAPsec* gene pair on both the X and the Y chromosome occurred independently several times in members of the *D. obscura* group, where this normally autosomal gene pair is sex-linked due to a sex chromosome - autosome fusion. Investigation of gene expression and short RNA profiles at the *S-Lap1*/*GAPsec* system suggest that meiotic drive and suppression likely involves RNAi mechanisms. Our finding suggests that recurrent conflict over sex chromosome transmission has shaped widespread genomic and evolutionary patterns, including the epigenetic regulation of sex chromosomes, the distribution of sex-biased genes, and the evolution of hybrid sterility.

## Introduction

Selfish genetic elements whose evolutionary trajectories are in conflict with those of their host were first described almost 100 years ago [1]. However, only in the recent decade has it become apparent that the antagonistic coevolution resulting from genetic conflict has shaped genome content and structure across the tree of life, from bacteria to plants and animals [2]. Antagonistic coevolution can occur between organisms, as in the evolutionary “arms race” experienced between pathogens and their hosts, or within genomes, where genetic elements can manipulate meiosis (or gametogenesis) so that they are transmitted to more than 50% of offspring (so called “segregation distorters” or “meiotic drivers”)[3]. These processes can leave behind a variety of distinct genetic signatures. For example, genes involved in pathogen virulence and host resistance consistently show elevated levels of amino acid substitutions (i.e. dN/dS) whereas genes involved in intra-genomic conflict, such as segregation distorters and their suppressors, often have high rates of lineage-specific duplications and gene amplifications (see below).

While segregation distorters can arise on all chromosomes, they are particularly prone to evolve on sex chromosomes [4]. Most characterized segregation distortion systems involve a distorting locus that targets a sensitive responder locus on its homolog, so that the chromosome where the driver resides is transmitted to more than 50% of the offspring of a heterozygous carrier [5]. A distorter must be able to discriminate its host chromosome from its homolog, and the distorter and responder loci must be in strong linkage to avoid the generation of suicide chromosomes that carry the distorter and a sensitive responder [4]; both of these conditions are met on differentiated, non-recombining X and Y chromosomes. The propensity of sex chromosomes to evolve distorting alleles could be strong enough to drive several unique characteristics of sex chromosomes, and it has been suggested that heterochromatinization of the Y and meiotic inactivation of the X during spermatogenesis evolved in part as protective measures to silence such alleles [5,6]. If this is the case, young sex chromosomes may be particularly susceptible to evolving sex ratio distortion because they often lack these features.

Disruption of equal transmission of the X and Y chromosomes will skew the population sex ratio and thus create strong selective pressure to evolve loci that repress sex chromosome drive [7,8]. Sex chromosome drive may thus be prevalent in evolution but is often cryptic, since bursts of meiotic drive elements arising on sex chromosomes should be followed by the quick invasion of suppressor alleles that restore equal sex ratios [4,7]. Cryptic drive systems are thus typically only revealed through detailed genetic manipulations or crosses between populations or species.

While cryptic drive systems cannot be observed directly, recurrent bursts of invading drivers and repressors may leave common footprints at patterns of genome evolution that are indicative of past meiotic drive. In particular, molecular characterization of meiotic drive systems in both mammals and fruit flies has found the following commonalities: In several instances, suppressors were found to share common sequences with the distorter itself [9-12]. This is the case for the Winters sex-ratio system within *D. simulans*, for instance. Here, two X-linked distorters, *Dox* and *MDox* (with *Dox* being derived from *MDox*), are both necessary for drive [9,10], and two autosomal genes (called *Nmy* and *Tmy*) that both originated from *Dox*/*MDox* suppress this drive [13]. Often, both the distorter and suppressor alleles are not only homologous to each other, but also amplified on both the X and Y chromosome [12,14,15]. A cryptic sex chromosome drive in mouse, for example, involves the convergent acquisition and amplification of the same gene families (*Slx*/*Sly*) on both the X and Y chromosome, and careful experimentation has shown that the co-amplified genes are in a co-evolutionary battle over sex chromosome transmission, whereby the X-and Y-linked copies of a gene family directly silence each other [11,12]. *Sly* knockdowns show female-biased sex ratios, while *Slx* deficiency causes a sex ratio distortion towards males. A similar mechanisms of cryptic segregation distortion has been implicated in the *Stellate*/*Suppressor of Stellate* (*Ste*/*Su*(*Ste*) system in *D. melanogaster*, where the expression of the X-linked gene *Ste* leads to the production of defective sperm, and *Su*(*Ste*), which is a multi-gene copy of *Ste* that moved to the Y-chromosome, silences *Ste* [16,17]. Thus, cryptic sex chromosome drive has repeatedly led to characteristic patterns of gene amplification of homologous genes on both the X and the Y chromosomes (**Figure 1A**).

**Figure 1.**
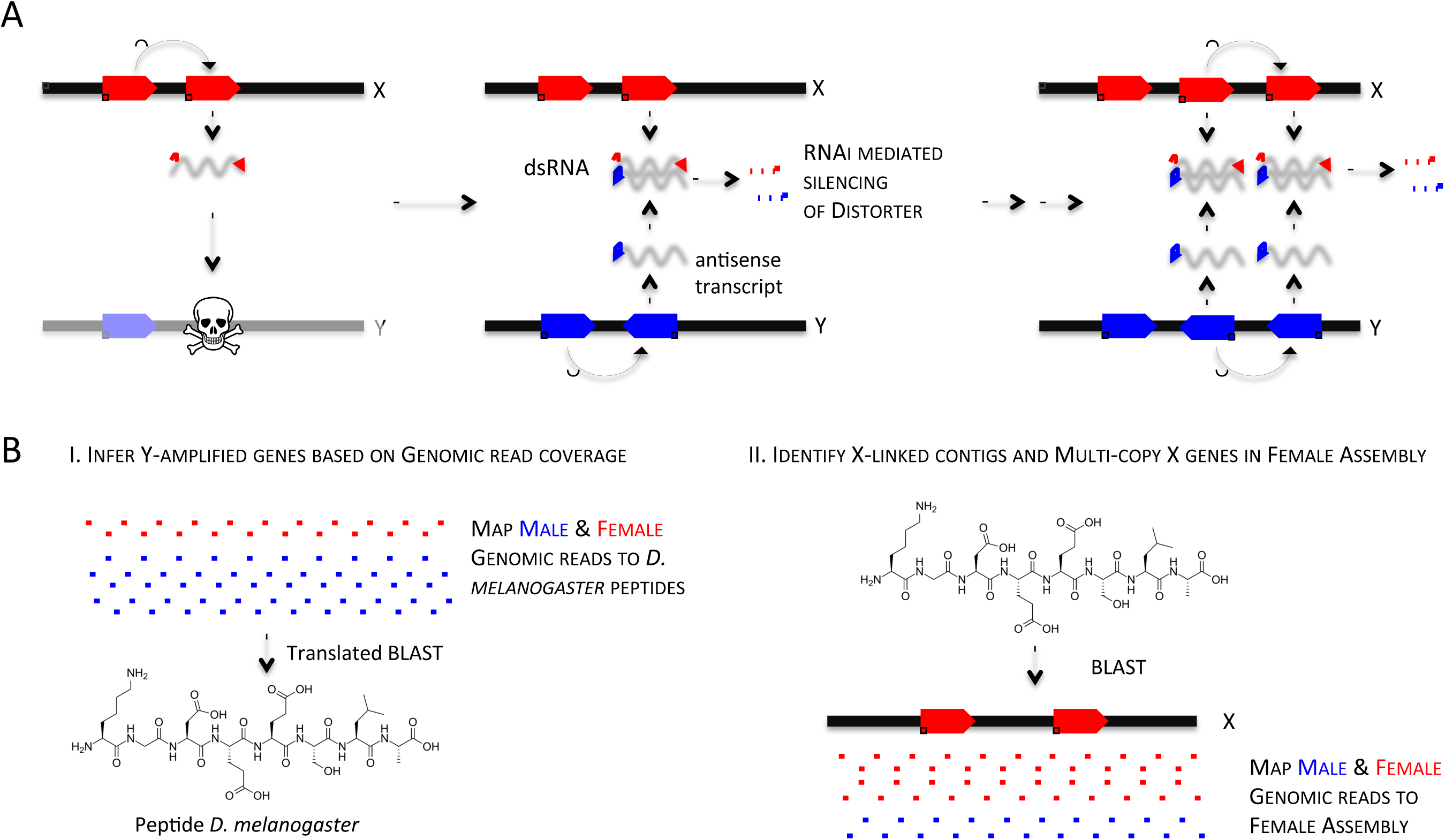
Meiotic conflict can fuel co-amplification of X and Y genes. **A.** An X-linked gene duplicate can evolve a novel function that eliminates Y bearing sperm. Amplification of the homologous Y gene, and production of antisense transcript may trigger the RNAi response, and silence the distorter. Repeated cycles of amplification of dosage-sensitive distorters and suppressors can result in the co-amplification of X/Y genes that are targeted by short RNAs. **B.** Bioinformatic identification of co-amplified X/Y genes. Multicopy Y genes are identified based on mapping of male and female genomic reads to D. melanogaster proteins using translated BLAST searches. Multi-copy X-linked homologs are identified for multi-copy Y genes, based on genome assemblies. Sex-linkage of contigs is inferred based on male and female read coverage of contigs, or based on published genome assemblies for a subset of species.

In several cases, suppression is likely mediated by sequence homology between the suppressor and distorter, through RNAi mechanisms resulting in posttranscriptional gene silencing. For example, piRNA pathway mutants enhance the drive caused by segregation distorter in *D. melanogaster* (the *Sd*/*Rsp* system) [18]. In addition, the sex-ratio driver Winters in *D. simulans* (*Dox*/*Nmy*) is suppressed through small RNA-based silencing [9,10,13], where *Nmy* and *Tmy* encode related hairpin RNAs that generate siRNA that repress their paralogous distorters [13]; knockout of RNAi pathway genes derepresses these sex-ratio drivers and disrupts spermatogenesis [13]. Similarly, the Y-linked *Su*(*Ste*) in *D. melanogaster* silences *Ste* through anti-sense expression and generation of small RNA’s [16]. This shows that the emergence of homologous repeated sequences can resolve an intragenomic conflict, through invoking the RNAi pathway.

Thus, cryptic sex chromosome drive has repeatedly led to characteristic patterns of gene amplification of homologous genes on both the X and the Y chromosomes that are targeted by short RNAs. Sex chromosomes also provide opportunity for other types of intragenomic conflict to arise. Sex chromosomes are hotspots for sexually antagonistic genes (i.e. genes that are beneficial to one sex but detrimental to the other [19,20]), and Y chromosomes often contain genes involved in male fertility that can undergo antagonistic coevolution due to sperm competition [21,22]. Recent studies have shown that the Y chromosomes of many organisms contain testes-specific genes that have amplified in copy number [23-25]. Some of these Y-linked gene families, such as those in mice, have been shown to be involved in sex chromosome drive, whereas for other gene families, the extra copies may either act to increase gene dosage or prevent degeneration by providing a substrate for non-allelic gene conversion [26]. Alternatively, the extra copies may be neutral or even slightly deleterious, yet they remain on the Y due to the reduced efficiency of selection on this non-recombining chromosome [27]. One key signature that appears to be unique to Y-amplified genes involved in sex chromosome drive is that their X-linked homologs have duplicated as well. This pattern is consistent with antagonistic coevolution resulting from repeated bouts of sex ratio distortion and suppression.

Here, we use bioinformatics and functional genomic analyses to assess the importance of sex chromosome drive across Drosophila species. Consistent with a role for Y-amplified genes unrelated to sex chromosome drive (see discussion), we find hundreds of genes that appear to be present in multiple copies on the Y chromosomes of various Drosophila species. However, we also find a second category of Y-amplified genes whose X homolog has been duplicated as well. We show that species with young sex chromosomes have repeatedly evolved genes that have co-amplified on the X and the Y, and show functions and expression patterns that are consistent with widespread meiotic conflict. Detailed investigation of one of these putative drive systems in the *pseudoobscura* group suggests that the putative driver is targeted by RNAi, and reveal that the same genes appear to have become involved in a meiotic conflict independently among multiple species of this group.

## Results

### Bioinformatic inference of co-amplified X and Y genes across Drosophila

To analyze gene content evolution and identify amplified X- and Y-linked genes, we sequenced both male and female genomic DNA in 26 Drosophila species from across the Drosophila phylogeny (**Table S1**). Roughly half of the species considered (11 out of 26) harbor the typical sex chromosome complement of Drosophila (that is, a single pair of ancient sex chromosomes, shared by all members of Drosophila). In addition to the ancestral pair of sex chromosomes, the other 15 species have a younger pair of “neo-sex” chromosomes, which formed when an autosome became fused to one or both of the ancient X and Y chromosomes (**Figure S1**). These younger “neo-sex” chromosomes are at various stages of evolving the typical properties of ancestral sex chromosomes, with neo-Y chromosomes losing their original genes and acquiring a genetically inert heterochromatic appearance, and neo-X chromosomes acquiring their unique gene content and sex-specific expression patterns [27,28]. We identified putative Y-amplified genes based on male and female gene coverage without relying on a genome assembly (**Figure 1B**, see Methods) and validated our approach using a high-quality genome assembly from *D. pseudoobscura* (**Figure S2**). Using this approach, we identify (depending on our cutoffs) 100s of genes that have multiple copies on the Y across the 26 species investigated (**Table S2, S3**, **Figure S3**). Genes might amplify on the Y for a variety of reasons, but co-amplification of testis genes may be a defining feature of genes evolving under meiotic conflict (see Discussion).

Among these multi-copy gene families on the Y, we found 46 amplified Y-linked genes with co-amplified X homologs in 10 species (**Table 1, Table S4**). We infer that the copy number of these co-amplified X/Y genes ranges from 8 copies on the Y up to 297 Y-linked copies (for an uncharacterized testis gene in *D. melanogaster* that amplified on the Y of *D. robusta*), with a mean copy number of 58 (see **Table S5**). We detect between 2-4 X-linked copies in our assemblies for these co-amplified X/Y genes (**Table S5**). However, the number of assembled X copies is likely an underestimate, since recent gene duplicates are typically collapsed in assemblies derived from short read sequencing data (see also Discussion). Several features of these co-amplified genes are typical of meiotic drive systems identified in flies and mammals. First, consistent with their proposed action of cheating fair meiosis, many are highly expressed in reproductive tissues in *D. melanogaster* (**Table S4**), and several of the genes have meiosis-related functions that may be exploited for sex chromosome drive (**Table 1, Table S4**). For example, we identify genes that are associated with spindle assembly involved in male meiosis (*fest*), chromosome segregation (*mars*), or male meiosis cytokinesis (*scra*), amongst others (**Table 1**).

**Table 1.**
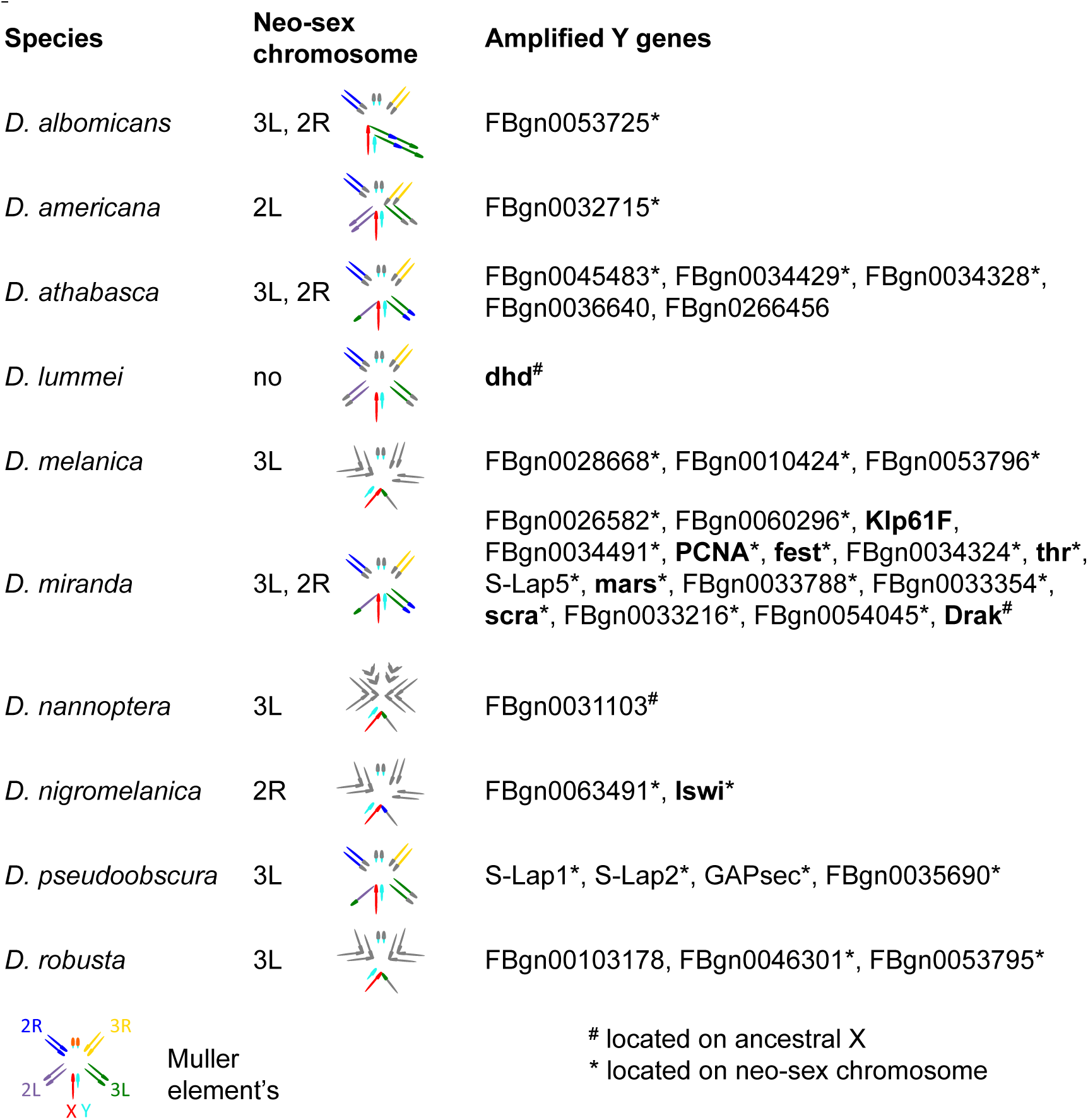
**Multi-copy Y-linked genes across Drosophila species**. Shown is which chromosome arms form the neo-sex chromosomes (based on synteny in *D. melanogaster*). Genes in bold have functions related to chromosome segregation.

Amplified Y genes were detected in each species investigated (**Table S2**). Interestingly, however, co-amplification of genes is much more common in species with recently added neo-sex chromosomes: of the 10 species where we found co-amplified genes, nine harbor neo-sex chromosomes (**Table 1**). Young sex chromosomes have many more genes that can evolve to cheat meiosis, and in the vast majority of cases the amplified genes were ancestrally present on the chromosome that formed the neo-sex chromosomes (**Table 1**). Also, young sex chromosomes may not yet be transcriptionally down-regulated during spermatogenesis, and sex-linked drivers are thus less likely to be silenced [5].

### Characterization of *S-Lap1* / *GAPsec* gene family in *D. pseudoobscura*

We decided to more carefully characterize a putative cryptic drive system in *D. pseudoobscura,* a species with a high quality PacBio-based genome assembly. *D. pseudoobscura* currently lacks an assembled Y chromosome, but we inferred Y-linkage of contigs based on male and female read coverage using Illumina data (see Methods). We identified two adjacent genes that exist in multiple copies on the X and Y chromosome of *D. pseudoobscura*: *S-Lap1* (Dpse\GA19547) and *GAPsec* (Dpse\GA28668). *S-Lap1* is a member of a leucyl aminopeptidase gene family that encodes the major protein constituents of Drosophila sperm [29], while *GAPsec* is a GTPase activating protein. This situation is reminiscent of the *Segregation distorter* meiotic drive system in *D. melanogaster*, where the distorter is a tandem duplication of RanGAP, which is also a GTPase activator [30]. Both *S-Lap1* and *GAPsec* show partial tandem duplications on the X (**Figure 2A**), and we detect roughly 100 (partial and full-length) copies of both *S-Lap1* and *GAPsec* on the Y chromosome (the Y-linked contigs contain 127 copies of *S-Lap1* and 91 copies of *GAPsec*; **Figure 2B, Figure 3**). *S-Lap1* appears to have independently duplicated in different parts of the Drosophila phylogeny (**Figure 2C**), but phylogenetic clustering of *S-Lap1* and *S-Lap2* in certain species groups could also result from gene conversion homogenizing gene duplicates within a clade [31]. In most species of the Drosophila clade, *S-Lap1* and *S-Lap2* are similar in size; in *D. pseudoobscura* and its sister *D. persimilis*, however, *S-Lap2* has acquired a large deletion, removing more than half of the 3’ end of the gene (**Figure 2A**; **Figure S4**). The partial duplication of *GAPsec*, on the other hand, is only found in *D. pseudoobscura* and its close relative *D. persimilis* (**Figure 2A, C**). *S-Lap1* and *GAPsec* probably dispersed onto the Y chromosome simultaneously, as there are multiple locations on the Y that preserve their X orientation (**Figure 2B**); note however, that the amplified copies on the Y do not include the tandemly duplicated copies. For both *S-Lap1* and *GAPsec*, the X- and Y-linked copies are highly expressed in testis of *D. pseudoobscura* (**Table S4, Figure 4, Table S6**), as expected for genes that try to cheat fair male meiosis.

**Figure 2.**
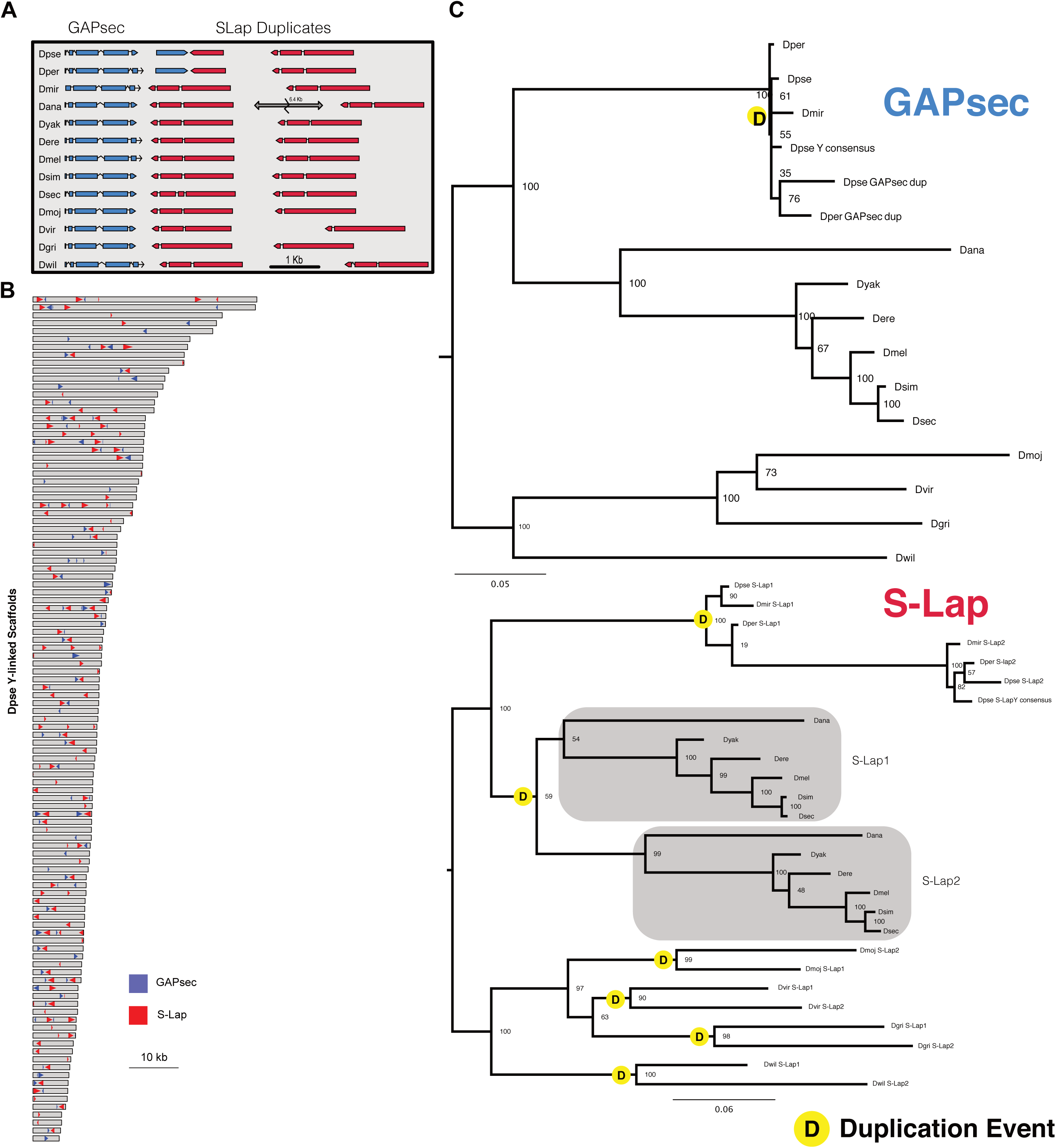
Molecular evolution of the *S-Lap1* and *GAPsec* genes in Drosophila. **A.** Organization of *S-Lap1* and *GAPsec* genes across the Drosophila genus. *S-Lap1* is duplicated in all Drosophila species investigated, and *GAPsec* shows a partial duplication in *D. pseudoobscura* and its sister species *D. persimilis*. **B.** Amplification of *S-Lap1* and *GAPsec* on Y-linked scaffolds of *D. pseudoobscura*. **C.** Gene trees of *S-Lap1* and *GAPsec* copies. *S-Lap1* independently duplicated multiple times across the Drosophila phylogeny. Inferred duplication events are shown by the yellow circle.

**Figure 3.**
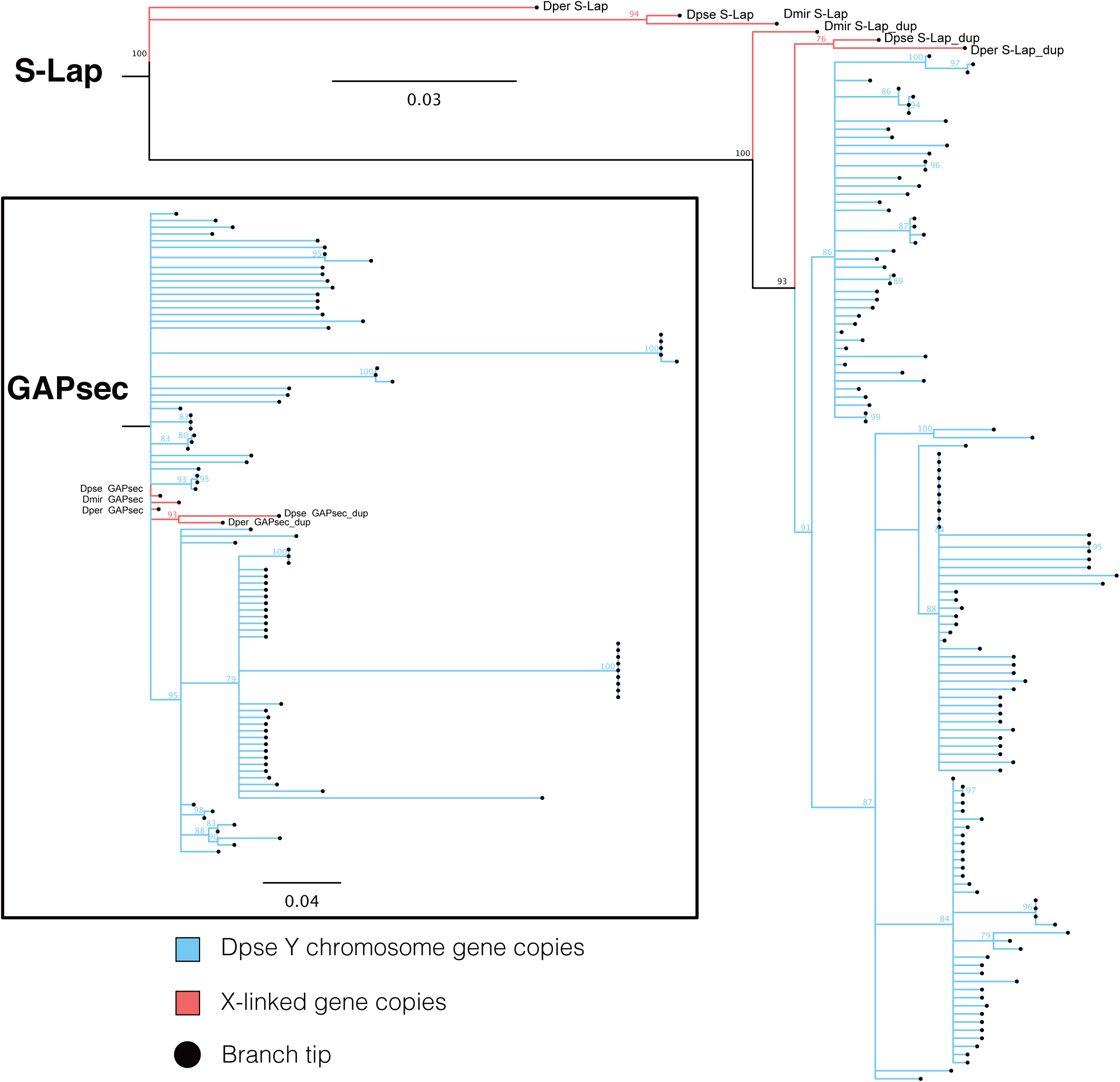
Phylogeny of *S-Lap1* and *GAPsec* gene families in *D. pseudoobscura*. Y-linked copies are shown by blue branches, and X-linked copies are shown in red.

In several cases, meiotic drive and suppression involves RNAi mechanisms [13]. We used stranded RNA-seq and small RNA profiles from wildtype *D. pseudoobscura* testes, to obtain insights into the molecular mechanism of the putative meiotic drive system involving *S-Lap1* and *GAPsec*. Interestingly, we detect both sense and antisense transcripts and short RNA’s derived from *S-Lap1* (**Figure 4A, Table S6;** see **Figure S5, S6** for the size distribution of short RNAs mapping and **Figure S7, S8** for cross-mapping of RNA-seq and short RNA reads between different gene copies). In particular, stranded RNA-seq data reveal that the X-linked copy of *S-Lap1*-duplicate produces both sense and anti-sense transcripts, resulting in the production of short RNA’s (see **Figure 4A**, **Table S6**). Close inspection of this genomic region in *D. pseudoobscura* shows that the duplicated *GAPsec* gene is directly adjacent to where the *S-Lap1*-duplicate antisense transcript begins (**Figure 4A**). Intriguingly, this segment scores highly as a potential promoter sequence when using the Berkeley Drosophila Genome Project (BDGP) neural network promoter prediction algorithm [32] (score=0.89, highest possible score=1). Thus, this suggests that the partial duplication of *GAPsec* provided a promoter-like sequence in *D. pseudoobscura* for antisense transcription of *S-Lap1*-duplicate. Note that this putative promoter sequence is not part of the Y copies of *S-Lap1* (which lack the *GAPsec* duplicate), and we detect virtually no antisense transcripts that originate from the Y-amplified copies of *S-Lap1* (**Table S6**). CAGE-seq data support that the *GAPsec* duplicate generated a new TSS resulting in antisense transcript of *S-Lap1*-dup (**Figure 4A**).

**Figure 4.**
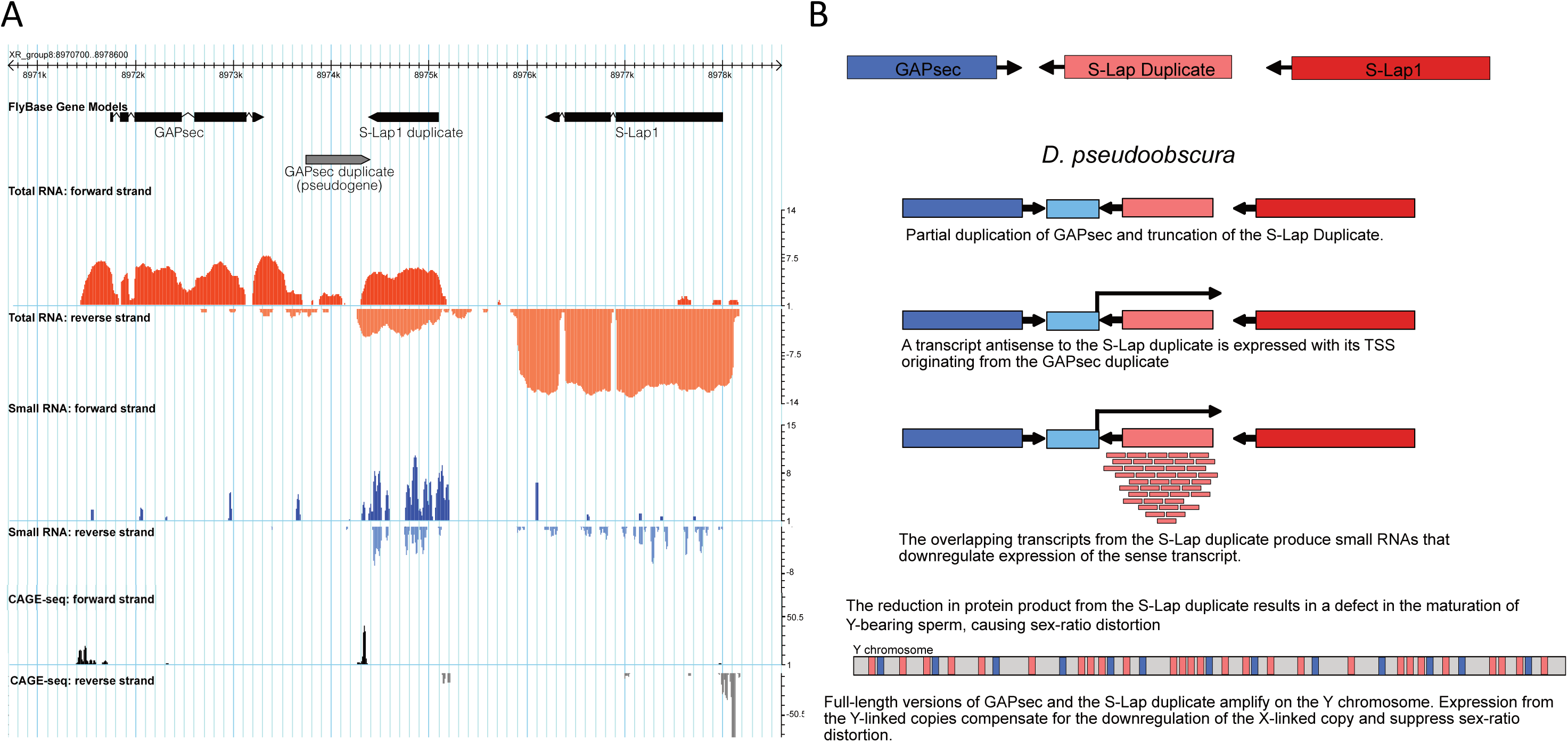
Short RNA’s may be involved in suppression of the cryptic *S-Lap1* drive system. **A.** Expression and short RNA profiles from wildtype *D. pseudoobscura* testis. Stranded RNA-seq (red tracks) reveals that the X-linked copy of *S-Lap1*-duplicate produces both sense and anti-sense transcripts, resulting in the production of short RNA’s (blue tracks). CAGE-seq data (grey tracks) support that the *GAPsec* duplicate generated a new TSS resulting in antisense transcript of *S-Lap1*-dup. **B.** A hypothetical evolutionary model of the cryptic *S-Lap1* drive system. *S-Lap1* was duplicated in an ancestor of *D. pseudoobscura*, and a partial duplication of *GAPsec* created a TSS for anti-sense transcription of *S-Lap1* duplicate. Production of small RNA’s may deplete *S-Lap1* transcripts, which may result in elimination of Y-bearing sperm, and could be compensated by amplification of *S-Lap1* on the Y chromosome.

These data suggest a possible model of cryptic sex chromosome drive in *D. pseudoobscura* (**Figure 4B**). *S-Lap1* is the most abundant sperm protein in *D. melanogaster* [29] but its function is unknown. If this protein is crucial for generating Y-bearing sperm, depletion of *S-Lap1* during spermatogenesis would result in drive. *S-Lap1* was duplicated in an ancestor of *D. pseudoobscura*, and a partial duplication of *GAPsec* (and truncation of *S-Lap1*-duplicate) created a TSS for anti-sense transcription of *S-Lap1* duplicate. Anti-sense production of *S-Lap1*-dup transcript may trigger siRNA production and silencing of *S-Lap1*, which could result in elimination of Y-bearing sperm. Acquisition of multiple copies of S*-Lap1* on the Y chromosome could restore *S-Lap1* function, and create a cryptic drive system in *D. pseudoobscura* (**Figure 4B**). Detailed molecular testing will be necessary to characterize the wildtype function of *S-Lap1*, and the cellular basis of the drive phenotype and its suppression.

### Independent co-amplification of *S-Lap1* and *GAPsec* in other flies of the *Drosophila obscura* group

In most Drosophila species, *S-Lap1* and *GAPsec* are located on an autosome (chromosome 3L in *D. melanogaster*). In the *D. pseudoobscura* and *affinis* group, however, this chromosome arm fused with the sex chromosomes about 15MY ago, causing *S-Lap1* to become sex-linked. Intriguingly, patterns of molecular evolution at *S-Lap1* and *GAPsec* suggest that they may be involved in meiotic conflicts in several members of the *pseudoobscura* species group. We used high-quality PacBio genome assemblies for two additional members of that species group [33], *D. miranda*, which diverged form *D. pseudoobscura* about 2-4 MY ago, and *D. athabasca*, which diverged 10-15 MY ago [34]. While our Illumina sequencing-based approach failed to detect co-amplified X and Y genes in these species (see **Table 1**), examination of the assembled PacBio genomes revealed that both gene pairs independently amplified on the sex chromosomes of both *D. miranda* and *D. athabasca* (see **Figure 5**). We identify tandem duplications of the entire genomic region containing a total of 11 copies of *S-Lap1* and 6 copies of *GAPsec* on chromosome XR in *D. miranda*, and these two genes have amplified 5 and 4 times, respectively, on the neo-Y chromosome of *D. miranda* (**Figure 5B**). Both the nature of the duplication event and patterns of sequence evolution suggest that co-amplification of *S-Lap1* and *GAPsec* occurred independently in *D. miranda*. Here, the XR copies arose from individual duplications of these two genes followed by three tandem duplications of the entire genomic region encompassing *S-Lap1* and *GAPsec*, producing a total of 11 copies of *S-Lap1* and 6 copies of *GAPsec*. All six of the X-linked copies of *GAPsec* are highly similar to each other (>99% identical), and more similar to their Y-linked paralogs than they are to *D. pseudoobscura* **(Figure 5A**). Also, *S-Lap1* and *GAPsec* appear to have moved only to a single location on the neo-Y of *D. miranda*, instead of being dispersed all across the Y, as in *D. pseudoobscura* (**Figure 5B**). Patterns of gene expression and short RNA production in *D. miranda* mimic that of *D. pseudoobscura*, with *SLap-1* (and *GAPsec*) transcripts being produced from both strands, and short RNA’s are generated across that genomic region (**Figure 6**). The mechanism of antisense production appears to differ from *D. pseudoobscura* (**Figure 6**). In particular, transcriptional read-through at both *S-Lap1* and *GAPsec* appear to generate anti-sense transcripts of both genes. Again, this would provide dsRNA targets for both *S-Lap1* and *GAPsec* in *D. miranda* that trigger the RNAi response, and select for copy number increases of those genes on the neo-Y, to silence the driving X. Close inspection of this genomic region in *D. miranda* reveals sequence differences between the X and Y copies that may account for antisense production at X-linked gene copies. In particular, we detect a polyadenylation signal (AATAAA) for *GAPsec* that is present in most (3 of the 4) Y copies, and in the homologous copy on the X in *D. pseudoobscura*, but which is missing in the *D. miranda* X-linked copies of *GAPsec*. This mutational event could account for the production of read-through transcripts on the X of *D. miranda*, leading to production of antisense transcript for *S-Lap1* and initiation of RNAi, analogous to the model proposed for *D. pseudoobscura*.

**Figure 5.**
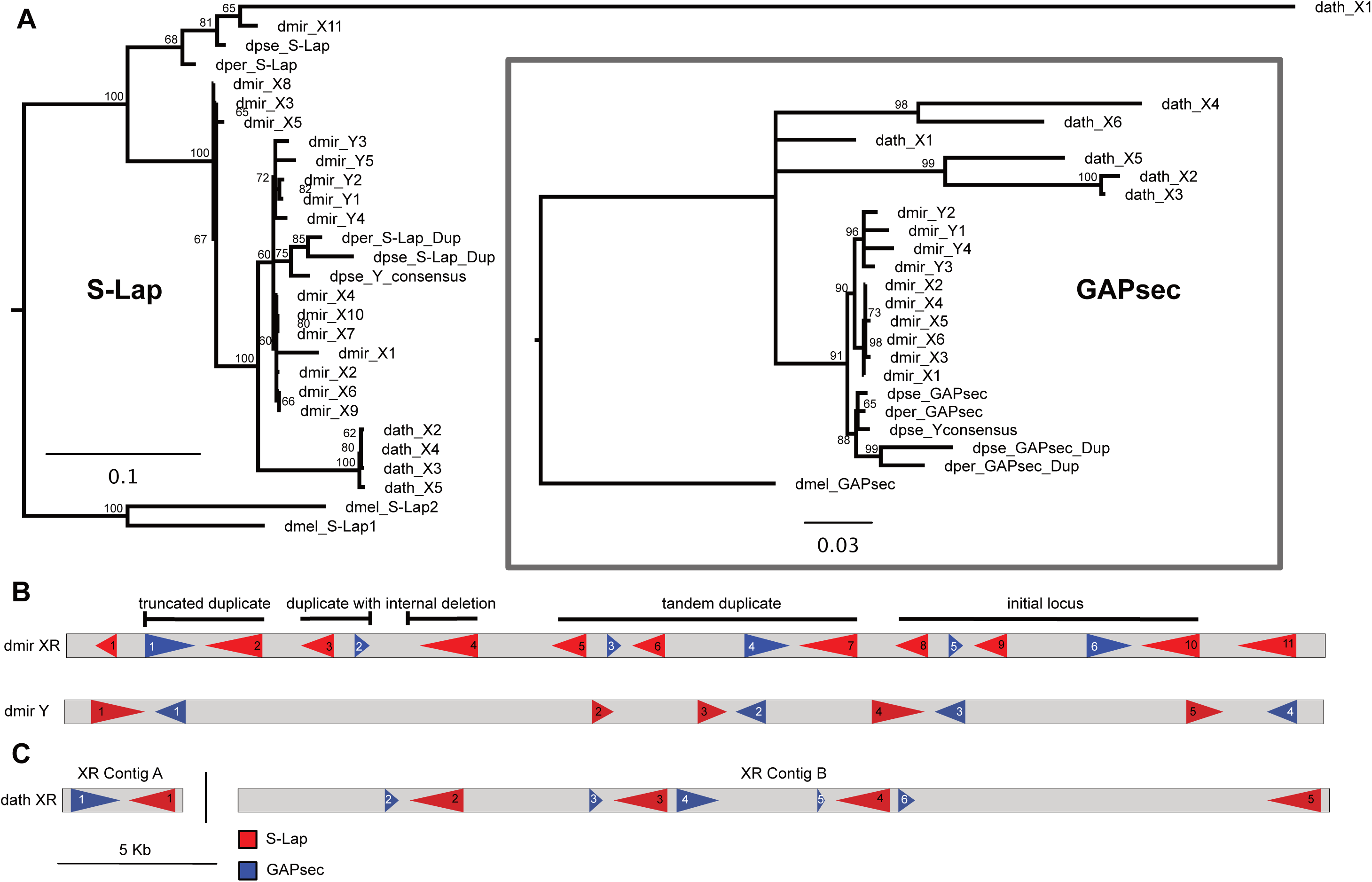
Independent amplification of S-Lap1 / GAPsec in different species of the *obscura* group.

**Figure 6.**
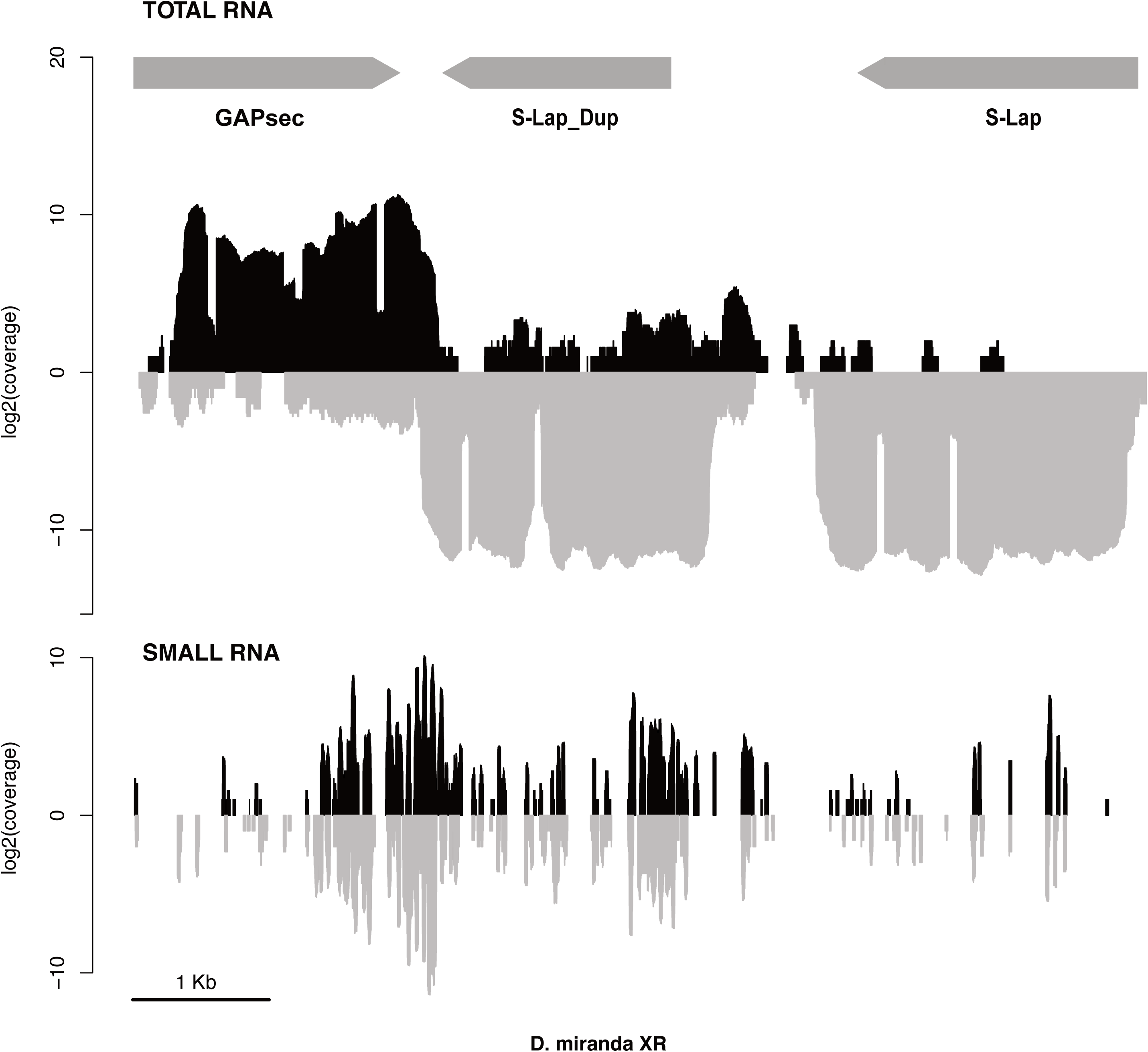
Short RNAs may be involved in suppression of the cryptic *S-Lap1* drive system in *D. miranda.* For legend, see Figure 4A.

Likewise, we infer independent amplifications of *S-Lap1* and *GAPsec* on the X chromosome of *D. athabasca*, another member of the *pseudoobscura* group for which we have generated a high-quality female genome assembly (**Figure 5C**). We detect 6 copies of *GAPsec* and 5 copies of *S-Lap1* on the X chromosome of *D. athabasca*, and genomic coverage analysis suggests a similar number of copies on the Y chromosome. This indicates that *S-Lap1* and *GAPsec* are involved in sex chromosomes drive several times in species where this locus has become sex-linked.

## Discussion

Theory predicts that selfish genetic elements that cheat fair meiosis may be common on sex chromosomes [4]. Sex chromosome drive has extreme evolutionary consequences, and distorted sex ratios can lead to population collapse and extinction, unless suppressors evolve rapidly to silence drivers on sex chromosomes [7]. Recurrent bursts of invasion of sex chromosome drivers followed by the quick fixation of repressors may thus be common, and may be a main evolutionary force driving the rapid evolution of sex determination mechanisms [35], shape the distribution of genes and patterns of gene expression on sex chromosomes [5], and lead to the evolution of hybrid sterility and account for the special role the X plays in the evolution of reproductive isolation [36,37]. However, given the short-lived nature of sex chromosome drive, the prevalence and the molecular bases of such conflicts remains poorly understood.

### Co-amplification of sex-linked genes is common in Drosophila

Here, we identify cryptic sex chromosome drive systems based on the footprints they leave behind in patterns of genome evolution: co-amplification of genes on both the X and the Y chromosome. Genes may amplify on the Y chromosome for a variety of reasons, and our current data do not allow us to evaluate their relative importance. In particular, multi-copy genes may simply arise on the Y at a higher rate, since the high repeat content on the Y facilitates structural re-arrangements that can promote gene family expansion [38]. Additionally, the efficacy of natural selection is reduced on the non-recombining Y, and Y chromosomes across diverse taxa accumulate functionless and deleterious repetitive DNA [27]. Amplified Y genes thus may either provide no benefit for their carriers, or could in fact be slightly deleterious, yet natural selection is unable to remove them [39]. Heterochromatin formation on the Y may further dampen any functional consequences of gene family expansion, and multi-copy Y genes may simply be more tolerated on the silenced Y. Finally, some multi-copy Y genes may actually contribute to male fitness and fertility [23-25,40]. Y chromosomes are transmitted from father to son, and are thus a perfect genomic location for genes that specifically enhance male fitness [41]. Y chromosomes of several species, including mammals and Drosophila, have been shown to contain multi-copy gene families that are expressed in testis and contribute to male fertility [23-25]. Our analysis shows that multi-copy Y genes are common across flies, and it will be of great interest to identify the diverse evolutionary processes driving their amplification.

Co-amplification of X/Y genes, on the other hand, is difficult to explain under scenarios that do not involve meiotic conflict: The repeat content of X chromosomes is comparable to that of autosomes [33]; natural selection efficiently purges deleterious mutations from the recombining X; and transcription of the X chromosome in Drosophila males is increased, rather than reduced [42]. Most importantly, co-amplified X and Y genes are enriched for meiosis functions (see also [43]), and the X-linked copies of co-amplified genes are highly expressed in testis [43]. Testis expression of co-amplified X-linked genes is unusual, as the X chromosome of Drosophila is normally devoid of testis-expressed genes [44], but can be understood under intragenomic conflict models invoking cryptic sex chromosome drive [5,9,10,45,46].

Our comparative analysis shows that co-amplification of genes on sex chromosomes is common in Drosophila and we find evidence of cryptic sex ratio distortion systems across distantly related species. Note, however, that our method for identifying co-amplified X/Y genes is conservative, and we might greatly underestimate the true magnitude of cryptic sex chromosome drive. On one hand, our approach for detecting amplified Y-genes requires them to have much higher coverage in male than female genomic reads (i.e. 4-fold higher coverage), and can thus only detect genes that have acquired considerably more copies on the Y chromosome relative to the X. Indeed, our recent careful examination of gene family evolution on the fully sequenced and assembled neo-Y of *D. miranda* confirms that the true number of co-amplified X/Y gene families is much higher than what we can detect here: Direct sequence inspection revealed that at least 94 genes co-amplified on the X and Y of *D. miranda* [43], while we could only detect 16 genes with our methodology. In addition, we only probed for genes that are present in the *D. melanogaster* annotation. Most of the species that we surveyed here are only distantly related to *D. melanogaster*, and genes that may have evolved novel drive-related functions may simply not have a homolog in *D. melanogaster.* Indeed, about 1/3 of the co-amplified X/Y genes that we identified in *D. miranda* did not have an ortholog in *D. melanogaster* [43]. Finally, we required X-linked co-amplified gene copies to be present in our Illumina assemblies; however, recent gene duplicates are often collapsed is such assemblies [33].

### Co-amplification of sex-linked genes and meiotic drive

Our data suggest that several putative drive genes have well-characterized functions in meiosis ensuring precise chromosome segregation, and selfish elements have recurrently succeeded to manipulate these normally tightly regulated cellular processes. How does co-amplification of meiosis-related genes on the X and Y cause meiotic drive and its suppression? If amplified Y genes are involved in a battle with the X over fair transmission, changes in gene copy number may tip the balance over inclusion into functional sperm, and could result in repeated co-amplification of distorters and suppressors on the sex chromosomes (**Figure 1A**). In particular, an X-linked gene involved in chromosome segregation may evolve a duplicate that acquires the ability to incapacitate Y-bearing sperm (**Figure 1A**). Invasion of this sex-ratio distorter skews the population sex ratio and creates a selective advantage to evolve a Y-linked suppressor that is resistant to the distorter. Suppression may be achieved at the molecular level by increased copy number of the wildtype function or by inactivation of X-linked drivers using RNAi [9,10,13]. If both driver and suppressor are dosage sensitive, they would undergo iterated cycles of expansion, resulting in rapid co-amplification of both driver and suppressor on the X and Y chromosome [4].

Co-amplification of X/Y genes is thus a characteristic signature of recurrent drive, but does not tell us which chromosome started the battle over inclusion into the sperm. That is, who drives whom? While we cannot determine the sequence of evolutionary events with certainty, the X chromosome is more likely to acquire segregation distorters, for several reasons. X chromosomes typically harbor many more genes that could acquire a driving function. Y chromosomes have a lower effective population size, and natural selection is impaired on the non-recombining Y [27], making drivers less likely to originate and spread on the Y. In addition, old Y chromosomes in many species are highly heterochromatic, and defects in heterochromatin formation can lead to chromosome mis-segregation [47]. This makes the Y chromosome especially vulnerable for mis-segregation, and the X chromosome—via DNA-binding proteins or other proteins involved in the regulation of the chromatin state—can take advantage of the heterochromatin state of the Y chromosome to induce meiotic drive. Indeed, most of the cellular meiotic drive phenotypes described in Drosophila species suggest a failure in chromatin state regulation [4,9,10,48-51]. We therefore expect *a priori* that most drivers originate on the X, and create strong selective pressure for the Y to evolve suppressors.

Most of the species where we identify co-amplified X/Y genes harbor neo-sex chromosomes. This suggests that sex ratio distorters have repeatedly evolved to exploit genomic vulnerabilities associated with the formation of new sex chromosomes. Different features of young vs. old sex chromosome create different susceptibilities to sex chromosome drive. Old Y chromosomes are typically highly repetitive and heterochromatic, a feature that may easily be exploited by a driver on the X. Also, old sex chromosomes show much higher levels of sequence divergence, which makes identification and targeting of the homolog by a driver easier. Yet, young Y chromosomes typically contain many more genes that can evolve to cheat meiosis, thereby increasing the chances of a Y-linked driver. Finally, young X chromosomes may not yet be transcriptionally inactive during spermatogenesis and thus express more drivers. In many species, including Drosophila, expression from the X chromosome is reduced during spermatogenesis [52]. As discussed above, the low gene number yet high repeat content makes Y chromosomes especially vulnerable to meiotic drive, and silencing of the X during spermatogenesis may have evolved as a genome defense against driving X’s [5]. Suppression of transcription during spermatogenesis may not yet have fully evolved on young X chromosomes, allowing the expression of more X-linked drivers.

### RNAi and drive

Our data further support growing evidence that the production of antisense transcripts, hairpin RNAs and small RNAs may be a common feature of meiotic drive elements [13,43]. RNA interference (RNAi) related pathways provide defense against viruses and transposable elements, and have been implicated in the suppression of meiotic drive elements [13]. Intriguingly, genes in these pathways often evolve rapidly, and show frequent gene duplication and loss over long evolutionary time periods. *Argonaute 2* (*Ago2*), for example, is one of the key RNAi genes in insects, and has repeatedly formed new testis-specific duplicates in the recent history of the *Drosophila obscura* group [53]. Analysis of additional RNAi-pathway genes confirms that they undergo frequent independent duplications and that their history has been particularly labile within the *Drosophila obscura* group [54]. Our finding suggests that the presence of young sex chromosomes in this species group makes them especially vulnerable to the invasion of meiotic drive elements, and may thus drive the rapid evolution of RNAi genes in this clade. It will be of interest to study the dynamics of RNAi genes in other species groups that have gained novel sex chromosomes, to see if diversification of RNAi genes is correlated with the emergence of new sex chromosomes.

In order to trigger the RNAi response, the production of dsRNA is required. This can be achieved in multiple ways. In the *D. simulans* Winters system, the two suppressor genes both encode related long inverted repeats that can form hairpin RNAs (hpRNAs), which are then processed by the RNAi machinery to generate siRNAs that repress the paralogous distorters [13]. Alternatively, the production of dsRNA can occur through anti-sense transcription of the target genes, and this mechanism is creating siRNAs in the putative drive involving *GAPsec* and *S-Lap1*, both in *D. pseudoobscura* and *D. miranda*.

The RNAi pathway can be utilized in different ways to either create a meiotic driver, or to suppress it. For example, a gene on the X (or it’s duplicate) may gain a novel function that disrupts segregation of the Y chromosome. The homologous Y gene (or duplicates of it) may then silence the driving X by producing anti-sense transcripts that generate dsRNAs and launch the RNAi response, to silence the X-linked driver. This scenario resembles the Winters sex ratio system, even though the suppressors of X-linked drive are autosomal and not Y-linked. The RNAi machinery can also be hijacked to create a driving X. In particular, if an X-linked gene is required for producing Y-bearing sperm, an X chromosome that silences this gene could evolve a drive phenotype. It could do so by antisense RNA production of this X-linked gene (or duplicates of it), thereby triggering the RNAi response to inactivate the gene. The organism could restore the wildtype function of this gene by increasing its dose through its amplification on the Y, and even non-functional copies may act as a decoy to soak up endo-siRNA that are targeting this locus. This pathway may underlay the putative *GAPsec* /*S-Lap1* drive system in *D. pseudoobscura*, where the X-linked duplicates of *S-Lap1* produce the vast majority of antisense transcripts (roughly 95%, see **Table S6**), while the Y-linked *S-Lap1* copies predominantly generate sense RNA (>99.9%). Most of the short RNA are produced from *S-Lap1-dup* (the putative driver) and the Y-linked copies of *S-Lap1* (about 96% in total), consistent with the idea that amplification of this spermatogenesis gene allows restoration of wildype function, possibly by acting as a decoy to dilute RNAi induced silencing triggered by antisense transcripts of *S-Lap1-dup.* Under either scenario, if both the driver and suppressor are dosage-sensitive, this can lead to the repeated invasion of driving and suppressing chromosomes through co-amplification of genes on the X and Y chromosome.

## Conclusion

To conclude, our comparative analysis suggests that cryptic sex chromosome drive may be common, especially on young sex chromosomes. The prevalence of cryptic sex ratio drive systems in insects and animals may account for several evolutionary and molecular phenomena [5]. Sex ratio distorters can fuel the rapid turn-over of sex determination mechanisms across species [35], and the transcriptional inactivation of sex chromosomes during spermatogenesis may have evolved as a defense against meiotic drive elements. The recurrent fixation of cryptic drive systems on sex chromosomes might explain the prominent role of the X chromosome in the evolution of hybrid sterility in a wide range of species [36,37,55-57], and contribute to genomic biases in the location of sex-biased genes or gene duplicates [58,59]. Our work suggests that the short RNA pathway has an important role not only in controlling genomic parasites such as transposable elements, but also selfish genetic elements that try to exploit highly regulated cellular proceeses such as chromosome segregation [60]. Future characterization of the putative drive systems identified here will provide a full picture of how distorting elements manipulate and cheat meiosis, what molecular pathways or developmental processes are particularly vulnerable, and how the genome has launched evolutionary responses to counter distortion.

## Methods

### Genome sequencing & assembly

Strains were acquired from the Drosophila Species Stock Center (UC San Diego) or the EHIME stock center (Ehime University, Japan) as indicated in **Table S1**. For each strain, DNA was extracted from a single male and a single female, using the Qiagen Gentra Puregene cell kit. The Illumina TruSeq Nano DNA library preparation kit was used to prepare 100 bp paired-end sequencing libraries for all species except *D. robusta*, *D. melanica*, and *D. willistoni*. For these species, the Illumina Nextera DNA library preparation kit was used to prepare 150 bp paired-end sequencing libraries. The genome assemblies produced for this study are noted in **Table S1**. Assemblies were produced from the female data: reads were error-corrected using BFC [61] and assembled using IDBA-UD [62] with default parameters.

### Identification of X-A fusions

X chromosome/autosome fusions were identified in two steps [63]. For each species, genomic scaffolds were assigned to Muller elements based on their gene content, inferred from the results of a translated BLAST search of *D. melanogaster* peptides to the assembly of interest. Scaffolds smaller than 5kb were excluded. Next, the male and female Illumina data were separately mapped to the female assembly using Bowtie2 [64] and excluding alignments with mapping quality less than 20. The coverage ratio (M/F) was calculated for each scaffold that was assigned to a Muller element. The distribution of coverage ratios for each Muller element (**Figure S1**) was then examined to determine if any of the ancestral autosomes had become X-linked.

### Identification of co-amplified genes on the X- and Y-chromosome

To characterize co-amplified genes on the sex chromosomes, we first identify genes amplified on the Y. For each species, male and female Illumina reads were separately aligned to a filtered version of the *D. melanogaster* peptide set, where only the longest isoform of each gene was retained. To generate these alignments, the DIAMOND software package [65] was used to perform a translated search of each Illumina read to the peptide set. Read coverage for each peptide sequence was calculated in 30 amino acid non-overlapping windows and normalized by dividing by the total number of mapped reads. The M/F coverage ratio was computed by dividing the median male coverage by the median female coverage, for each peptide. We required that potentially Y-amplified genes have a normalized M/F coverage ratio of at least 2.5 and only retained genes whose parent copy was X-linked in the species of interest. We searched for X-linked duplicates in the female genome assemblies by first using Exonerate [66] to extract the coding sequence of the best hit between the *D. melanogaster* peptide and the female assembly. We then used BLASTN [67] to obtain a stringent (E-value threshold = 1e-20) list of all non-overlapping hits between each exon of the coding sequence and the genome assembly. We considered a gene to be duplicated in females if at least 25% of the parent coding sequence aligned to more than one location in the genome assembly.

### *S-Lap1* and *GAPsec* gene trees and Y chromosome gene copies

The Muller-D copies of *S-Lap1* and *GAPsec* were identified in the 12 Drosophila genomes [68] by synteny with *D. melanogaster* and their coding sequences were downloaded from FlyBase [69]. The PRANK software package [70] was used to generate codon-aware alignments of coding sequences for each gene. The resulting alignment was trimmed using trimAl [71] and RaxML [72] was used to infer a maximum likelihood phylogeny (100 bootstrap replicates). *D. pseudoobscura* Y-linked contigs were identified using read coverage information from male versus female genomic sequencing data. Exonerate [66] was used to determine the location of the amplified copies of *S-Lap1* and *GAPsec* on these scaffolds with the *D. pseudoobscura S-Lap1* (*FBpp0285960*) and *GAPsec* (*FBpp0308917*) peptide sequences as queries. The *D. pseudoobscura* Y copies of each gene were aligned using MAFFT [73], trimmed with trimAl [71], and a Y consensus sequence for each gene was generated using PILER [74].

### RNA libraries and mapping

We dissected testes from 3-8 day old virgin males of *D. pseudoobscura* (strain MV25, transgenic *P{w^+^:sh-S-Lap1-Y}* and *white* control males) reared at 18°C on Bloomington food. We used Trizol (Invitrogen) and GlycoBlue (Invitrogen) to extract and isolate total RNA. *D. pseudoobscura* CAGE-seq data were obtained from the ModEncode project [75]. We resolved 20 μg of total RNA on a 15% TBE-Urea gel (Invitrogen) and size selected 19-29 nt long RNA, and used Illumina’s TruSeq Small RNA Library Preparation Kit to prepare small RNA libraries, which were sequenced on an Illumina HiSeq 4000 at 50 nt read length (single-end)., We used to Ribo-Zero to deplete ribosomal RNA from total RNA, and used Illumina’s TruSeq Stranded Total RNA Library Preparation Kit to prepare stranded testis RNA libraries, which were sequenced on an Illumina HiSeq 4000 at 100 nt read length (paired-end). Total RNA data were aligned to the *D. pseudoobscura* reference genome using HISAT2 [76], whereas Bowtie2 [64] (seed length: 18) was used to align small RNA and CAGE-seq data. In all cases, alignments with mapping quality less than 20 were discarded.

## Acknowledgements

Funded by NIH grants (R01GM076007, GM101255 and R01GM093182) to DB. We thank Lauren Gibilisco for generating short RNA libraries and testis RNA-seq libraries.

## Author contributions

C.E. generated the genome data and performed the bioinformatics analysis and D.B. oversaw the project and wrote the manuscript with input from all authors.

## Figure Legends

**Figure S1. Identification of newly formed sex chromosomes across Drosophila.** Sex chromosomes are inferred using male and female coverage data. Plotted is the normalized male / female genomic read coverage for scaffolds mapped to the *D. melanogaster* genome, to infer the location of Muller elements.

**Figure S2. Validation of bioinformatics pipeline to infer multi-copy Y genes in *D. pseudoobscura***. Shown is the predicted coverage based on mapping of Illumina reads on the x-axis, versus the number of Y-linked copies of a gene found in the genome assembly. Note that our bioinformatics pipeline is conservative and underestimates the number of Y-linked copies found in the assembly, presumably due to many multi-copy genes being fragmented in the assembly.

**Figure S3. Chromosomal location of multi-copy Y genes.**

**Figure S4. Alignment of X-linked and Y-linked copies of A. *S-Lap1* and B. *GAPsec***.

**Figure S5. Size distribution of short RNAs mapping to X and Y linked copies of *S-Lap1***.

**Figure S6. Size distribution of short RNAs mapping to X and Y linked copies of *S-Lap1***.

**Figure S7. Mappability of RNA-seq and short RNA data to X-linked and Y-linked copies of *S-Lap1*. FigureS8. Mappability of RNA-seq and short RNA data to X-linked and Y-linked copies of *GAPsec.***

**Table S1. Species used in this study**. Shown are species and stock numbers, total assembly size, and sex chromosome karyotype (see Figure S1).

**Table S2. Number of amplified Y genes**. Shown are the number of inferred amplified Y-linked genes found in each species, for a cut-off of male/female coverage ratio (M/F) >= 2.5.

**Table S3. Number of amplified Y genes identified, for different cut-offs of male/female coverage ratio (M/F from 2.5 to 10).**

**Table S4. Multi-copy Y-linked genes across Drosophila species**. Shown are the orthologous location of multi-copy Y genes in *D. melanogaster*, and their inferred molecular function and gene expression pattern in *D. melanogaster* (data from flybase.org).

**Table S5. Inferred copy numbers for co-amplified X and Y genes.**

**Table S6. Mapping of total RNA and short RNA for X-linked and Y-linked copies of *GAPsec* and *S-Lap1*.**

